# Carbon dioxide regulates *Mycobacterium tuberculosis* PhoPR signaling and virulence

**DOI:** 10.1101/2022.04.12.488064

**Authors:** Shelby J. Dechow, Rajni Goyal, Benjamin K. Johnson, Robert B. Abramovitch

**Affiliations:** Department of Microbiology and Molecular Genetics, Michigan State University, East Lansing, MI 48824

## Abstract

The *Mycobacterium tuberculosis* (Mtb) two-component regulatory system, PhoPR, is implicated in pH-sensing within the macrophage because it is strongly induced by acidic pH both *in vitro* and the macrophage phagosome. The carbonic anhydrase (CA) inhibitor ethoxzolamide (ETZ) inhibits PhoPR signaling supporting the hypothesis that CO_2_ may also play a role in regulating PhoPR. Here, we show that increasing CO_2_ concentration induces PhoPR signaling, and its induction is independent of medium pH. We also show that at acidic pH 5.7, a normally strong inducer of PhoPR signaling, that increasing CO_2_ from 0.5% to 5% further induces the pathway. Based on these findings, we propose that PhoPR functions as a CO_2_ sensor. Mtb has three CA (CanA, CanB, and CanC) and using CRISPR interference knockdowns and gene deletion mutants, we assessed which CAs regulate PhoPR signaling and virulence. We first examined if CA played a role in Mtb pathogenesis and observed that CanB was required for virulence in macrophages, where the knockdown strain had ~1 log reduction in virulence. To further define the interplay of CO_2_ and Mtb signaling, we conducted transcriptional profiling experiments at varying pH and CO_2_ concentrations. As hypothesized, we observed the induction of PhoPR at acidic pH is dependent on CO_2_ concentration, with a subset of core PhoPR regulon genes dependent on both 5% CO_2_ and acidic pH for their induction, including expression of the ESX-1 secretion system. Transcriptional profiling also revealed core CO_2_ responsive genes that were differentially expressed independently of the PhoPR regulon or the acidic pH-inducible regulon. Notably, genes regulated by a second two component regulatory system, TrcRS, are associated with adaptation to changes in CO_2_.

## Introduction

*Mycobacterium tuberculosis* (Mtb) virulence is dependent on its ability to sense environmental stimuli and adjust its physiology accordingly. One of the major intracellular stresses that Mtb faces is fluctuation in pH of the acidifying macrophage phagosome[1, 2]. The Mtb two-component regulatory system (TCS) PhoPR, is associated with Mtb pH sensing and slowing Mtb growth[1, 3, 4]. Over half of the PhoPR regulon is significantly up-regulated within two hours following macrophage infection, and its induction is dependent on phagosome acidification[1]. PhoPR is required for Mtb virulence in macrophages, mice, and guinea pigs, where deletion mutants are attenuated for growth in these models[5, 6]. PhoPR also controls sulfolipid expression[7, 8], which was recently shown to play a role in promoting cough and presumably transmission[9]. Thus, PhoPR could play a role for the duration of infection, from the initial stages of macrophage infection, survival and replication in macrophages and transmission to new hosts. While the PhoPR regulon is regulated by acidic pH, it is possible that it is directly or indirectly regulated by pH or possibly other reported signals like magnesium or chloride[6, 10]. Notably, the PhoPR regulon is inhibited by treatment with ethoxzolamide (ETZ)[11–13], a carbonic anhydrase (CA) inhibitor[14, 15], providing a potential link between carbon dioxide (CO_2_), pH sensing, and PhoPR regulation.

CO_2_ is a gas that plays a vital role in altering physiological and pathophysiological processes across all life, including photosynthesis, oxidative metabolism, and cell signaling[16]. As such, most organisms have evolved CO_2_-sensing mechanisms to adjust their physiology accordingly, implying that the ability to sense CO_2_ levels is key for organism survival. For bacteria, sensing changes in CO_2_ concentration is important for infecting and colonizing host tissues. Many bacterial species sense and respond to environmental shifts from ambient CO_2_ levels (0.03%) to higher CO_2_ levels (5%) as they enter their host organisms, initiating pathogenic differentiation[16, 17]. For example, *Vibrio cholerae* naturally inhabits aquatic ecosystems where it forms commensal or symbiotic relationships with marine organisms[18]. However, removal of pathogenic *V. cholerae* from aquatic environments and introduction into the human host induces virulence. The increase in CO_2_ levels found within the human host leads to subsequent increases in enterotoxin production in *V. cholerae* [19]. Specifically, *V. cholerae* relies on CA activity to initiate enterotoxin production and virulence is reduced following treatment with ETZ [20]. In *Mycobacterium bovis* BCG, CO_2_ was shown to play an important role in the growth under microaerophilic conditions and regulate a subset of *acr* co-regulated genes [21]. Therefore, CO_2_ sensing and CA activity may also play a role in Mtb virulence.

Carbonic anhydrases are ubiquitous metalloenzymes found in most biological organisms. These enzymes catalyze the essential interconversion of carbon dioxide (CO_2_), bicarbonate (HCO_3_^-^), and a proton (H^+^), a process that is characterized by rapid equilibration of all three components by CA[17]. Because CO_2_, HCO_3_^-^, and pH/H^+^ are in tight equilibrium with each other, and fluctuation in any one of these molecules can be reflected in the other two, pH can act as an indirect indicator of CO_2_ levels. For example, the low pH of gastric juices activates urea transport in *Helicobacter pylori*, resulting in high urease activity and CO_2_ production. *H. pylori* buffers periplasmic pH by relying on the conversion of CO_2_ to HCO_3_ via CA activity[22, 23], providing an example of indirect CO_2_ sensing by maintaining pH homeostasis. Mtb encodes for three of these carbonic anhydrases: Rv1284 (*canA*), Rv3588c (*canB*), and Rv3273 (*canC*). Based on global phenotypic profiling with transposon mutants, two of these CA (*canA* and *canB*) are predicted to be required for virulence in mice, suggesting that CO_2_-associated physiologies or sensing may be important for Mtb virulence[24].

ETZ is a potent inhibitor (~27 nM) of the most active recombinant CA protein in Mtb, CanB, and shows inhibitory activity in the low micromolar (~1.03 μM) and submicromolar (0.594 μM) range for recombinant CanA and CanC, respectively[14, 25, 26]. Our lab has previously confirmed that ETZ fully inhibits Mtb CA activity within cells, while also inhibiting the PhoPR regulon[11]. This suggests that a physiological link exists between CA activity and PhoPR signaling, and we hypothesize that ETZ may indirectly inhibit the PhoPR regulon by disrupting CA activity. Previously, we proposed a model where the interconversion of CO_2_ into HCO_3_ and a H^+^ may promote acidification of the Mtb pseudoperiplasm leading to activation of the PhoPR regulon[11]. ETZ would effectively block this process, disrputing PhoPR signaling. The goal of this study is to define interactions between CO_2_ concentrations, pH, CA, and PhoPR-dependent gene regulation and identify their functions in macrophage virulence.

## Materials and Methods

### Bacterial Culture Conditions

Experiments were performed with *M. tuberculosis* strain CDC1551, unless otherwise stated. Mtb was maintained in vented T-25 culture flasks in 7H9 Middlebrook medium supplemented with 10% oleic acid-albumin-dextrose-catalase (OADC), 0.05% Tween-80, and 0.2% glycerol and incubated at 37 °C with 5% CO_2_, unless noted otherwise. For experiments requiring buffered medium, 100 mM 3-(N-morpholino)propanesulfonic acid (MOPS) or 100 mM 2-(N-morpholino)ethanesulfonic acid (MES) was added to 7H9 medium for buffering to pH 7.0 or pH 5.7, respectively. Cultures were grown to mid-late log phase (OD_600_ 0.5-1.0) for use in experiments described below.

### Flow cytometry and fluorescence analysis

For flow cytometry experiments, *M. tuberculosis* CDC1551 (*aprA*’::GFP) was grown to mid-late log phase (OD_600_ 0.6-1.0) in non-inducing 7H9 medium buffered to pH 7.0. Cultures were pelleted, resuspended, and seeded at an initial OD_600_ of 0.2 into 8 mL of either non-inducing medium (7H9 [pH 7.0]) or GFP-inducing medium (7H9 [pH 5.7]). High (15%), medium (5%), or low (0.5%) CO_2_ concentration was applied to biological replicates of each culture condition. Cultures were incubated for six days after which samples were pelleted and fixed with 4% PFA. GFP fluorescence was measured using methods previously described by Abramovitch *et. al*. [3]

### Transcriptional profiling and data analysis

High-throughput RNA sequencing (RNA-seq) experiments were performed with Mtb CDC1551. Cultures were seeded at a starting OD_600_ of 0.2 in 8 mL of 7H9 buffered media and grown at 37°C in standing T-25 culture flasks. Two biological replicates of the following culture conditions were examined: i) 0.5% CO_2_ at pH 5.7, ii) 5% CO_2_ at pH 5.7, iii) 0.5% CO_2_ at pH 7.0, iv) 5% CO_2_ at pH 7.0. Cultures were incubated for six days, after which total bacterial RNA was extracted as previously described [1]. The SPARTA (ver. 1.0) software package was used to analyze raw sequencing data[27]. Differentially expressed genes were determined to have a differential gene expression > 1.5-fold and filtered based on log2CPM < 5. Gene enrichment was performed for Figure 4C and S4 using the hypergeometric distribution to determine statistical significance of gene overlap. Enrichment analysis for Figure 5A and 5B was performed using a Chi-Square analysis with Yates Correction. RNA-seq data has been deposited at the GEO database (Accession GSE200125).

### Construction of carbonic anhydrase CRISPRi knockdowns and ORBIT knockout

To investigate the role of carbonic anhydrases in Mtb pathogenesis, we inhibited expression of *Rv1284* and *Rv3588c* using the dCas9_Sth1_ CRISPRi system[28]. Single guide RNAs (sgRNAs) were designed with 20-22 nucleotides of complementarity to the target carbonic anhydrase (Table S1). *canC* could not be effectively knockeddown by CRISPRi, therefore, we generated a Δ*canC* knockout mutant using the ORBIT (oligonucleotide-mediated recombineering followed by Bxb1 integrase targeting) system and replaced *canC* with a hygromycin (Hyg^R^) resistance cassette. The ORBIT recombineering plasmid, pKM444, was electroporated into Mtb, and anhydrotetracycline (ATc) was added to induce expression of the RecT annealase and Bxb1 integrase. Electrocompetent cells were made from the pKM444 transformants and subsequently electroporated with the knockout integration plasmid, pKM464, and the *canC* targeting oligonucleotide. The *canC* knockout mutant was confirmed by sequencing the 5’ and 3’ junction sites using ORBIT target-specific and *canC*-specific primers (Table S1). Semi-quantitative, real-time reverse transcription PCR (qRT-PCR) was used to confirm loss of functional *canC*. Electrocompetent Δ*canC* was used to transform the CRISPRi *canA* and *canB* and generate a knockdown or knockout of all three Mtb carbonic anhydrases.

### Macrophage infections

Bone Marrow-derived macrophages (BMDMs) were extracted from mouse femurs and tibiae and cultivated at 37 °C with 5% CO_2_ in 24-well tissue culture plates as previously described[29]. BMDMs were infected at a multiplicity of infection (MOI) of 1:1 with the panel of CDC1551 CRISPRi strains unless otherwise stated. Fresh media was exchanged every two days and infected BMDMs were exposed to the following treatment conditions: a) bone marrow macrophage medium [BMMO], b) BMMO + 250 ng/μL Anhydrotetracycline (Atc), c) BMMO + 100 μM Ethoxzolamide (ETZ), and d) BMMO + 250 ng/μL Atc + 100 μM ETZ. Infected BMDMs were lysed by 0.1% v/v Tween 80 in distilled deionized water. Intracellular bacterial lysates were plated for days 0, 3, 6, and 9. Lysates were serially diluted and enumerated on 7H10 + 10% OADC agar plates and counted following 21 days of incubation at 37 °C. Each strain was performed in triplicate at the indicated timepoints.

### qRT-PCR

CRISPRi knockdown strains were incubated at ambient CO_2_ and 5%. CO_2_ levels with or without 250 ng/mL ATc and/or 100 μM ETZ. After six days of treatment, total RNA was extracted as previously described[1]. cDNA was generated using 1 μg of DNAase-treated RNA with the High-Capacity cDNA Reverse Transcription Kit (Applied Biosystems) and kit protocol. A reaction mix of 2 μL cDNA, 2 μL of forward and reverse qRT-PCR primer, 4 μL of DNAase-free H_2_O, and 10 μL of Power SYBR Green PCR Master Mix (Applied Biosystems) was made for each sample tested. All experiments were performed with two biological replicates separated into three technical replicates. The Quantstudio3 was used to perform the following qRT-PCR reaction: 95°C for 2 minutes followed by forty 2-step cycles of 95°C for 15 s and 60°C for 30 s. All samples were normalized to *sigA* signal and quantified using the ΔΔCT calculation.

## Results

### Carbon dioxide modulates the phoPR pathway independent of medium pH

The discovery of ETZ as an inhibitor of Mtb carbonic anhydrase activity and the PhoPR regulon suggested a potential link between CO_2_ and PhoPR signaling[11]. CO_2_ interacts with water to form carbonic acid (H_2_CO_3_), which quickly dissociates into a proton (H^+^) and bicarbonate (HCO_3_^-^). Therefore, when CO_2_ levels rise it causes a decrease in pH. We hypothesized that if we modulated CO_2_ concentrations, much like how Mtb experiences differences in in CO_2_ in the environment as compared to the lung, we may observe a subsequent modulation of the PhoPR regulon if it is indeed sensing the proton from CO_2_ dissolution. To investigate this, PhoPR signaling was monitored using the CDC1551 (*aprA*’::GFP) reporter strain in GFP-inducing (7H9, pH 5.7) and non-inducing (7H9, pH 7.0) media in ambient CO_2_ and 5% CO_2_ concentrations. We also treated flasks with 40 μM ETZ or DMSO. Notably, the media are strongly buffered with 100 mM MOPS or MES (pH 7.0 or 5.7 respectively), and changes in CO_2_ have no impact on the pH of the extracellular medium (as measured by pH color strips and pH-meter[11]). Cultures were incubated for six days, GFP fluorescence was normalized to optical density (OD), and samples were analyzed using a plate reader. We found that fluorescence of *aprA* at ambient CO_2_ in pH 5.7 medium with DMSO was ~1.5-fold lower compared to 5% (Figure 1A). Interestingly, ETZ causes an overall reduction in *aprA* fluorescence at pH 5.7; however, 5% CO_2_ does cause slightly higher fluorescence (Figure 1A). This observation is consistent with the disruption of PhoPR signaling by ETZ and hints that higher CO_2_ concentration may overcome some of the inhibitory activity.

**Figure 1.**
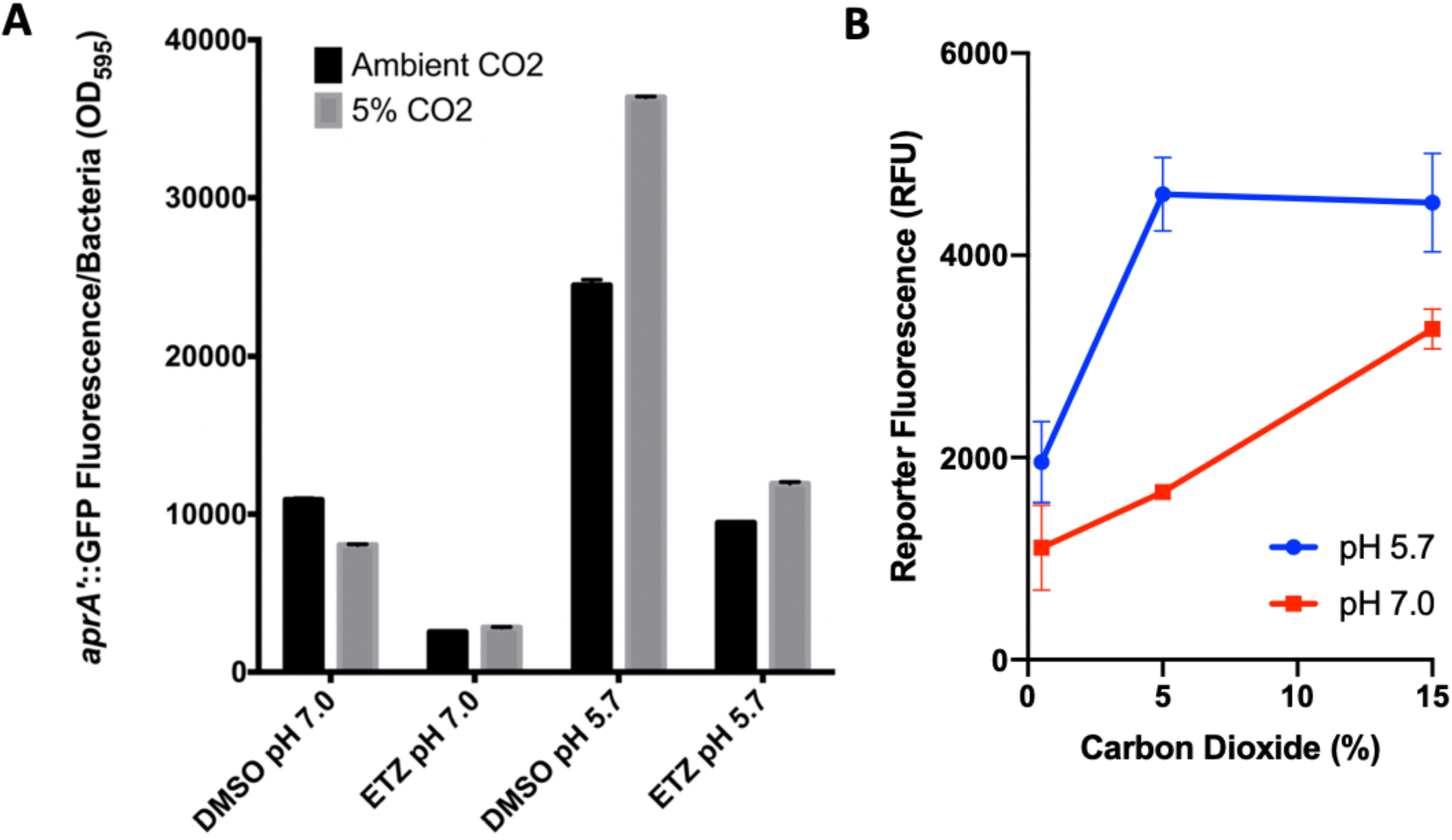
Changes in CO_2_ concentration directly modulate *phoPR*-regulated gene expression independent of extracellular pH. **A)** PhoPR-dependent CDC1551(*aprA*’::GFP) fluorescent reporter is responsive to changes in environmental carbon dioxide. CDC1551(*aprA*’::GFP) was grown in 7H9 rich medium buffered to pH 5.7 or 7.0 and exposed to ambient or 5% CO_2_ for six days. Conditions were performed in duplicate. The error bars represent the standard deviation. **B)** PhoPR-regulated *aprA* is modulated by CO_2_ independent of pH. CDC1551(*aprA*’::GFP) was grown in 7H9 rich medium buffered to pH 5.7 or 7.0 and exposed to high (15%), medium (5%), or low (0.5%) carbon dioxide concentrations for six days. Conditions were performed in duplicate and results are representative of two independent experiments. The error bars represent the standard deviation.

To further analyze the impact of CO_2_ concentration on PhoPR signaling, we repeated the experiment using a glovebox with well controlled levels of 0.5%, 5%, and 15% CO_2_ in medium buffered at pH 5.7 or 7.0 with 100 mM MES or MOPS, respectively. To address potential impacts of CO_2_ on growth that could impact normalized readings on a plate reader, we analyzed fluorescence of individual cells using flow cytometry. In all these culture conditions, the pH of the medium did not change due to the high levels of 100 mM buffer in the media. Following six days of incubation, exposure to 5% CO_2_ at pH 5.7 resulted in significant induction of PhoPR reporter fluorescence compared to 0.5% CO_2_ (Figure 1B). This level of induction was maintained at 15% CO_2_ at pH 5.7. Similarly, PhoPR reporter fluorescence was induced at neutral pH by increasing CO_2_ concentrations. As in our previous studies, with CO_2_ at 5%, we observed the strong pH-dependent induction of the reporter. These finding reveal that PhoPR can be regulated independent of the pH of the medium and is in fact responsive to CO_2_ concentrations, a finding consistent with PhoPR acting as a CO_2_ sensor.

### CanB is essential for survival in macrophages

The Mtb genome encodes three annotated carbonic anhydrases, Rv1284 (CanA), Rv3588c (CanB), and Rv3273 (CanC) of which CanA and CanB are required for virulence in mice using global transposon mutant analyses[30]. Biochemical studies show that CanB has the highest catalytic activity of all three carbonic anhydrases and that ETZ inhibits CanB activity (K_I_= 27 nM)[14]. We hypothesized that ETZ is targets CanB, subsequently downregulating the PhoPR regulon, driving the previously described inhibition of Mtb growth in infected macrophages and mice treated with ETZ[11]. To further investigate the function of CanA, CanB, and CanC during infection, we generated CRISPRi knockdowns of *canA, canB*, and *canAB* and a knockout mutant of *canC* to achieve disruption strains of all three CA. Successful anhydrotetracycline (ATc)-induced CRISPRi knockdown in WT CDC1551 background was confirmed through qRT-PCR (Supplemental Figure 1), with approximately 10-fold and 7-fold reduction of *canA* and *canB*, respectively. CRISPRi of *canC* was not observed, despite attempts with multiple different CRISPRi constructs, so we generated a Δ*canC* knockout strain in WT CDC1551 using the ORBIT system[31]. Δ*canC* was then confirmed by sequencing the 5’ and 3’ junction sites, PCR amplification of the knockout region, and qRT-PCR (Supplemental Figure 2). The CRISPRi strains were introduced into WT CDC1551 and the Δ*canC* strain to achieve different combinations of *canABC* functional disruption and were confirmed with qRT-PCR (Supplemental Figure 2). Bone marrow derived macrophages (BMDMs) were infected initially with the WT CDC1551 CRISPRi panel (CRISPRi-*canA*, CRISPRi-*canB*, CRISPRi-*canAB*). The empty CRISPRi vector, pLJR965, was also electroporated into WT CDC1551 and used to infect BMDMs. The infected macrophages were treated with either ATc, ETZ, both ATc and ETZ, or had no treatment applied. At the end of a 9-day macrophage survival assay, we observed ~0.25-0.5-log decrease in growth in all strains treated only with ETZ (Figure 2A-E). In the CRISPRi-EV and CRISPRi-*canA*, we observe inhibition of growth by ETZ treatment, but no impact of ATc treatment, suggesting a limited role of *canA* in Mtb virulence in macrophages. In contrast, when ATc is applied to infected cells containing CRISPRi-*canB* and CRISPRi-*canAB*, we observe ~ 1-log decrease in bacterial growth (Figure 2C-E). This indicates that CanB is required for Mtb virulence in macrophages. Notably, there were no CFU differences in ATc-only treated CRISPRi-*canB* and ATc+ETZ-treated CRISPRi-*canB*, an observation consistent with CanB potentially being the target of ETZ. We also examined the role of CanC using the knockout strain. Following a 9-day macrophage survival assay, we observed similar ~0.5-log decrease in Mtb growth in strains treated only with ETZ, but no impact on virulence in cells missing *canC* (Figure 2F). Notably, in the Δ*canC*-CRISPRi-*canB* and Δ*canC*-CRISPRi-*canAB* we observed a significant reduction of virulence, with a loss of activity for ETZ treatment. Together, these data support that CanB is required for virulence in macrophages and that ETZ activity may be driven by targeting CanB.

**Figure 2.**
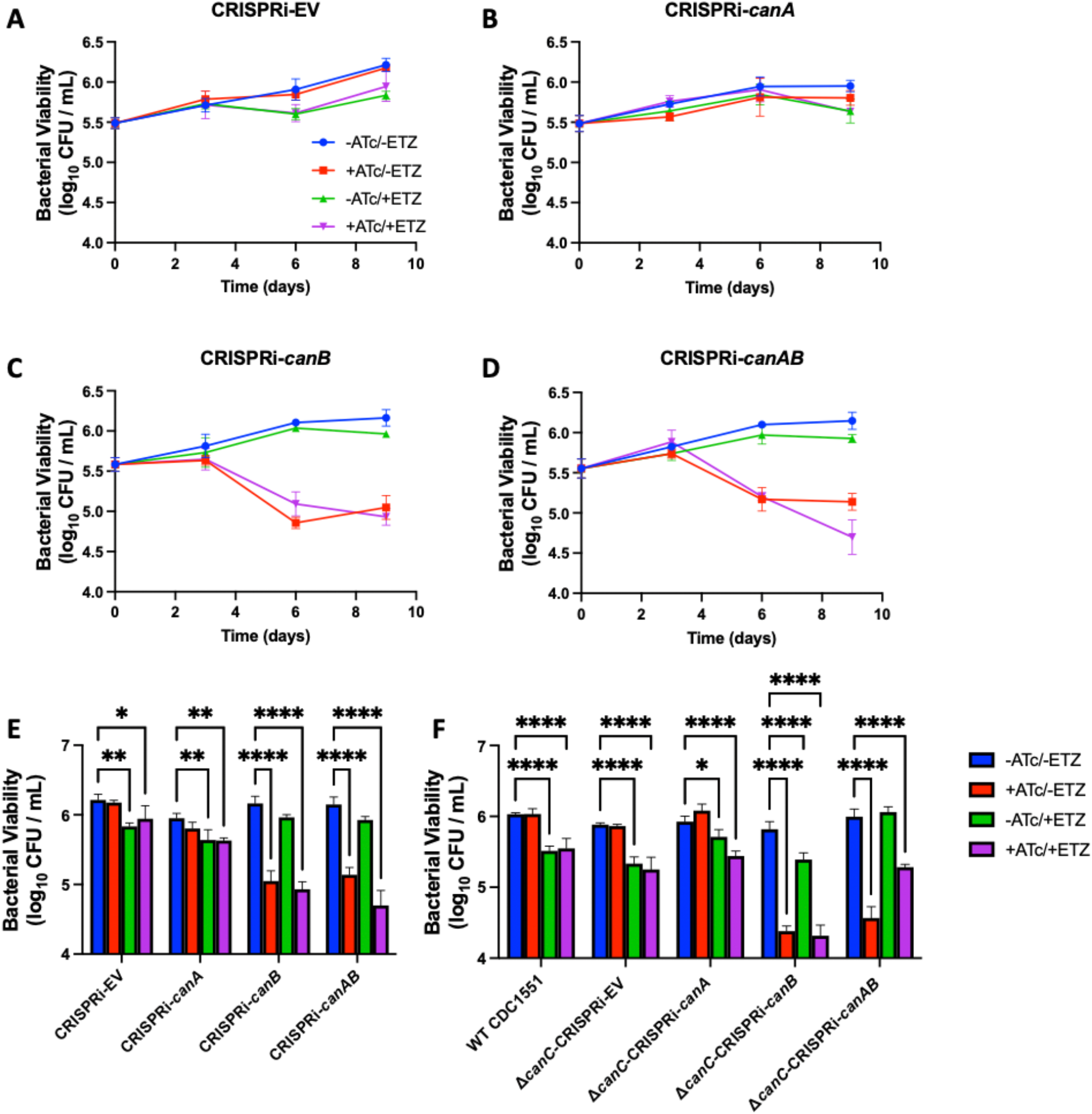
CRISPRi-*canB* exhibits reduced survival in macrophages. BMDMs infected with the CRISPRi strains in WT CDC1551: **A)** CRISPRi-EV, **B)** CRISPRi-*canA*, and **C)** CRISPRi-*canB*, and **D)** CRISPRi-*canAB*. All strain treatments were performed in triplicate over the course of 9 days and representative of multiple independent experiments. Error bars indicate standard deviation. Statistical analysis of growth differences between CRISPRi strains in the **E)** WT CDC1551 background and the **F)** Δ*canC* background at Day 9 in resting BMDMs. Significance was determined by one-way ANOVA (Tukey’s multiple comparisons test; *P < 0.05, **P < 0.01, ***P < 0.001, ****P < 0.0001). Mean ± SD are shown in the bar graph.

#### *canB* expression is not associated with changes in *aprA* expression

Based on our finding that *aprA* fluorescence is lower when incubated with ETZ at both pH 7.0 and pH 5.7 and that CanB is essential for pathogenesis, we examined *aprA* expression levels as a functional readout of PhoPR regulation, between ETZ treated Mtb and the *canB* knockdown. To test the hypothesis that CanB is required for PhoPR signaling, we incubated CDC1551 CRISPRi-EV and CRISPRi-*canB* in rich media buffered to pH 5.7 or pH 7.0 at 5% CO_2_ levels for six days in the presence of either 250 ng/mL ATc or 100 μM ETZ, both ATc and ETZ, or no treatment. *aprA* gene expression under each treatment condition for each strain was quantified by qRT-PCR relative to the no treatment CRISPRi-EV control. Contrary to our hypothesis, we did not observe repression of *aprA* gene expression when *canB* was knocked down with ATc treatment at either pH 5.7 or pH 7.0 (Figure 3A and B). *aprA* gene expression was, however, repressed ~10-fold when ETZ treatment was applied, which is consistent with the previous reporter fluorescence data and published RNAseq data[11] (Figure 1A). We investigated whether *canB* was appropriately knocked down by ATc. Indeed, we observed ~100-fold repression of *canB* transcript in the CRISPRi-*canB* treated with ATc (Figure 3C) compared to CRISPRi-*canB* with no treatment.

**Figure 3.**
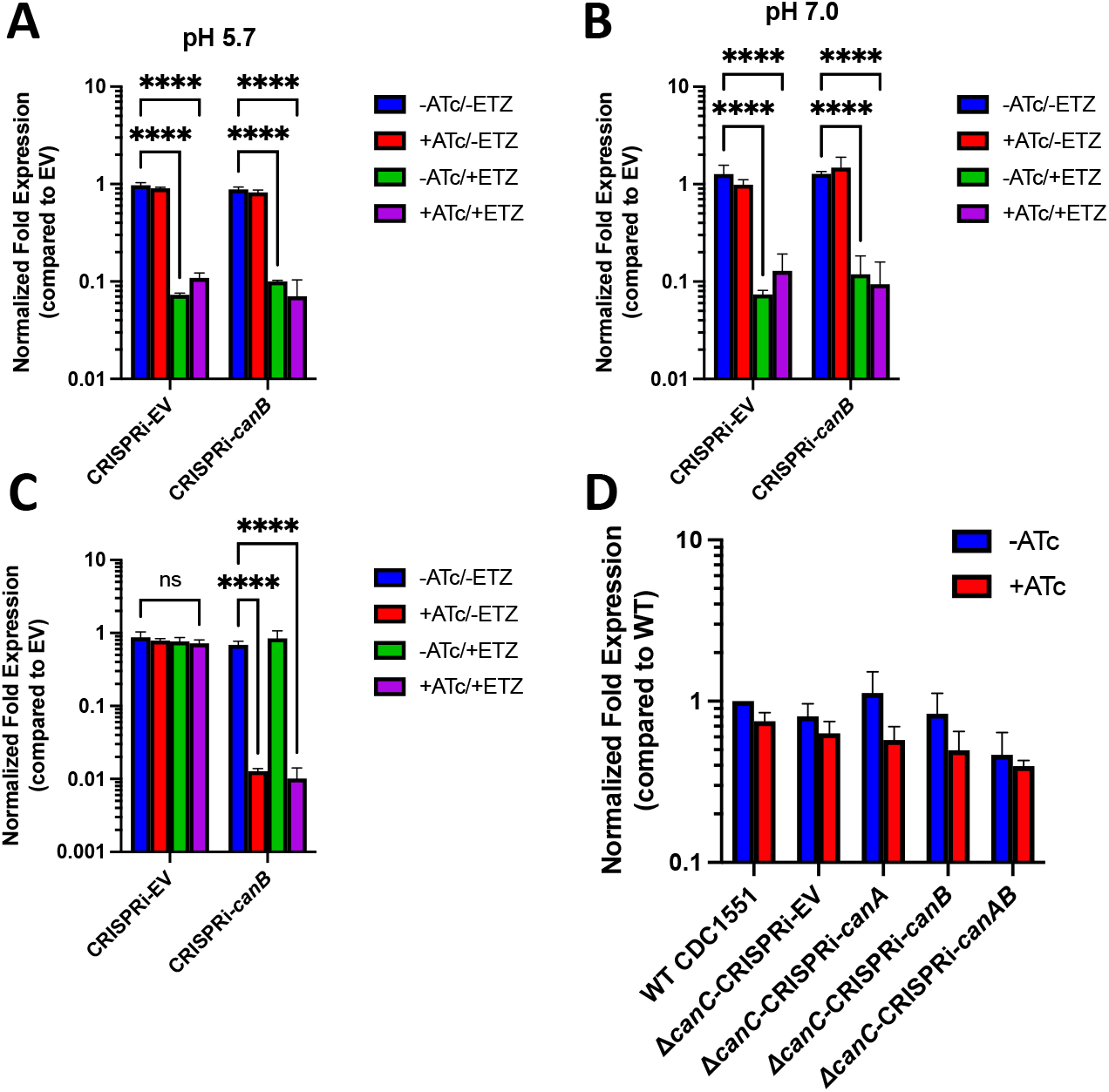
*aprA* expression is repressed in a CA-independent, ETZ-dependent manner. qRT-PCR comparing *aprA* expression of CRISPRi-EV and CRISPRi-*canB* in the WT CDC1551 background treated with either ATc, ETZ, both ATc and ETZ, or no treatment at **A)** pH 5.7 or **B)** pH 7.0 relative to the CRISPR-EV strain with no treatment. Data are shown as mean ± SD of three replicates. Statistical significance was determined using a two-way ANOVA (Tukey’s multiple comparisons test; ****P < 0.0001). **C)** qRT-PCR confirmation of *canB* expression knocked down in CRISPRi-*canB* when ATc treatment was applied compared to CRISPRi-EV. Data are shown as mean ± SD of three replicates. Two-way ANOVA was applied (Tukey’s multiple comparisons test; ****P < 0.0001, ns, not significant). **D)** qRT-PCR of *aprA* expression at pH 5.7 with Δ*canC* CRISPRi panel confirms that *aprA* is not repressed in a triple knockdown/knockout strain of Mtb CA. Data are shown as mean ± SD of three replicates.

In addition to *canB*, Mtb encodes for two additional CA genes (Rv1284 and Rv3273). We hypothesized that simultaneous knock down or knock out of all three CAs would lead to inhibition of *aprA* expression. To test this, we incubated WT CDC1551, Δ*canC*-CRISPRi-EV, Δ*canC*-CRISPRi-*canA*, Δ*canC*-CRISPRi-*canB*, and Δ*canC*-CRISPRi-*canAB*, in 7H9 buffered to pH 5.7 in the presence of absence of 250 ng/mL ATc for six days. *aprA* gene expression was quantified for each strain and treatment condition relative to the no treatment WT CDC1551 EV control. Again, we did not observe repression of *aprA* expression in double or triple knockdown and knockout strains of CA (Figure 3D). These data do not support our hypothesis that CA activity is directly modulating PhoPR signaling and suggest that ETZ inhibits PhoPR signaling *in vitro* by a mechanism that is independent of *canA*, *canB* or *canC* or that CA activity is not sufficiently disrupted by CRISPRi to modulate *aprA* expression.

### Genes induced by CO_2_ share significant overlap with the phoPR-regulon

Because increasing CO_2_ concentrations at pH 5.7 led to induction of the PhoPR-regulated *aprA* promoter, we investigated the extent to which the PhoPR regulon might be regulated by CO_2_. To define the CO_2_ transcriptional response in Mtb, we performed RNA-seq at pH 7.0 and pH 5.7 at 0.5% and 5%, CO_2_ and compared the transcriptional profiles (Table S2A-S2E). Mtb grown at pH 5.7 and exposed to 5% vs. 0.5% CO_2_ revealed global transcriptional changes with 183 genes induced (>1.5-fold, q<0.05) and 146 genes repressed (>1.5-fold, q<0.05) (Figure 4A). The majority of differentially expressed genes were involved in cell wall and cell processes and intermediary metabolism and respiration (Figure 4B), implying that the mycobacteria may be responding to and redirecting their metabolic activity due to the dynamic shift in CO_2_ concentration. In addition to *aprA* induction, we also observed a total of 39 induced genes that overlapped with genes from previously published data that have PhoP and acidic-pH dependent induction [11] (Figures 4C and 4D). Gene enrichment analysis of both transcriptional profiles revealed statistically significant overlap between the two groups of genes (p<0.0001). Notably, we observed widespread changes in gene expression in ESAT-6 secretion system-1 (ESX-1) protein secretion (*esxA, esxB, espABCDE, espL*). PE35 and PPE68 are located directly upstream of EsxA and EsxB and are required for Mtb virulence and EsxA and EsxB secretion[30, 32–34], were also differentially expressed in both profiles. ESX-1 secretion is regulated by PhoPR, and disruption of PhoP is known to negatively impact ESAT-6 secretion[6, 11, 35, 36]. The significant induction of ESX-1 associated genes in the 5% vs 0.5% CO_2_ transcriptional profile and significant repression in the *phoP::Tn* profile at pH 5.7 (Figure 4D), supports that this ESX-1 virulence activity is regulated by CO_2_ in a PhoPR dependent manner.

**Figure 4.**
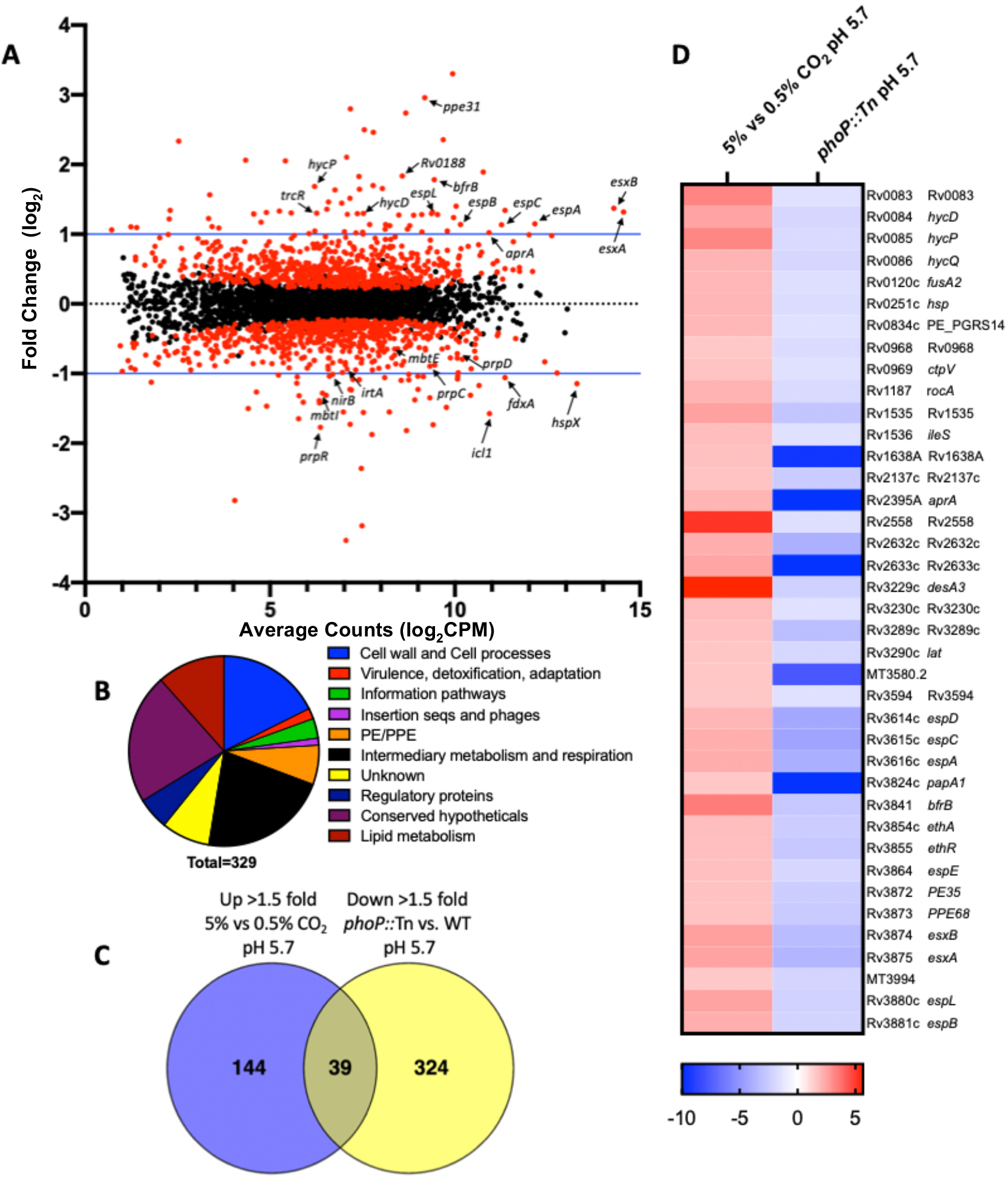
Increased CO_2_ concentration induces PhoPR-regulated genes at acidic pH. **A)** Mtb RNAseq transcriptional profiling data following exposure to 5% CO_2_ compared to 0.5% CO_2_ at pH 5.7. Upregulated genes that are indicated include PhoPR-regulated ESX-1 genes, hypoxia responsive genes, and the *hyc* locus. Downregulated genes that are indicated include those involved in iron acquisition and the methylcitrate cycle. Red dots denote statistically significant genes (q < 0.05). **B)** Pie chart depicting the functional categories of significantly differentially expressed genes (>1.5-fold, q < 0.05) derived from the pH 5.7, 5% CO_2_-treated Mtb RNA-seq transcriptional profile. **C)** Significant gene overlap observed between genes upregulated (Up) (>1.5-fold, q < 0.05) by 5% CO_2_ treatment at pH 5.7 and downregulated (Down) (>1.5-fold, q < 0.05) in the *phoP::Tn* mutant strain at pH 5.7[11]. **D)** A heat map of the overlapping 39 CO_2_-induced (red) and the *phoP::*Tn-repressed genes (blue) (>1.5-fold, q < 0.05). Genes are annotated using the H37Rv genome.

Downregulated genes in the 5% vs 0.5% CO_2_ transcriptional profile at pH 5.7 include those involved predominately in iron homeostasis and intermediary metabolism and respiration (Figure 4A). When comparing the downregulated CO_2_ profile to upregulated genes in the *phoP::Tn* profile, we see 26 differentially expressed genes that overlap (Supplemental Figure 4). Gene enrichment shows this overlap is statistically significant (p<0.0001). These overlapping genes include the iron-scavenging mycobactins (*mbtABCDEFGKL*) and the carboxymycobactin ABC transporter (*irtB*). In contrast, the iron storage gene (*bfrB*) is upregulated in the CO_2_ transcriptional profile and downregulated in *phoP::Tn* (Figure 4A), indicating that PhoPR regulation by CO_2_ impacts iron homeostasis. Together, roughly 20% of genes differentially regulated at 5% vs 0.5% CO_2_ at pH 5.7 are also PhoPR regulated (Figure 4C and D, and Supplemental Figure 4), strongly suggesting that the increased CO_2_ levels are inducing PhoPR signaling.

### RNA-seq studies define the CO_2_ regulon and implicate a role for TrcRS in responding to CO_2_

To define genes regulated by CO_2_ independent or dependent on pH, we initially compared genes regulated by CO_2_ (5% CO_2_ vs 0.5% CO_2_) at pH 7.0 or pH 5.7 and observed widespread changes at pH 7.0 with 78 genes induced (>1.5-fold, q<0.05) and 169 genes repressed (>1.5-fold, q<0.05) and at pH 5.7 with 183 genes induced (>1.5-fold, q<0.05) and 146 genes repressed (>1.5-fold, q<0.05) (Figure 4A).

We then specifically looked for CO_2_-regulated genes independent of pH regulation. In doing so, we compared the up-regulated transcriptional profiles of 5% CO_2_ vs 0.5% CO_2_ at both pH 7.0 and pH 5.7 with the acidic pH up-regulated and down-regulated genes (>1.5-fold, q<0.05) described in a previous RNA-seq study (that was conducted at 5% CO_2_)[11]. Forty-three genes are both acidic pH and CO_2_-upregulated of which 20 genes are controlled by PhoP including ESX-1 secretion genes (*esxAB, espABCDE*) and *aprA* (Figure 5 and Table 5). In the 24 genes downregulated by acidic pH and CO_2_, we see genes that are repressed by high iron conditions (*mbtBCF*) or involved in intermediary metabolism (*leuCD, pfkB*) (Figure 5B and Table 5). As previously reported under microaerophilic conditions and differential CO_2_[21], we also observed the downregulation of *hspX, acg*, Rv1733 and Rv1738.

**Figure 5.**
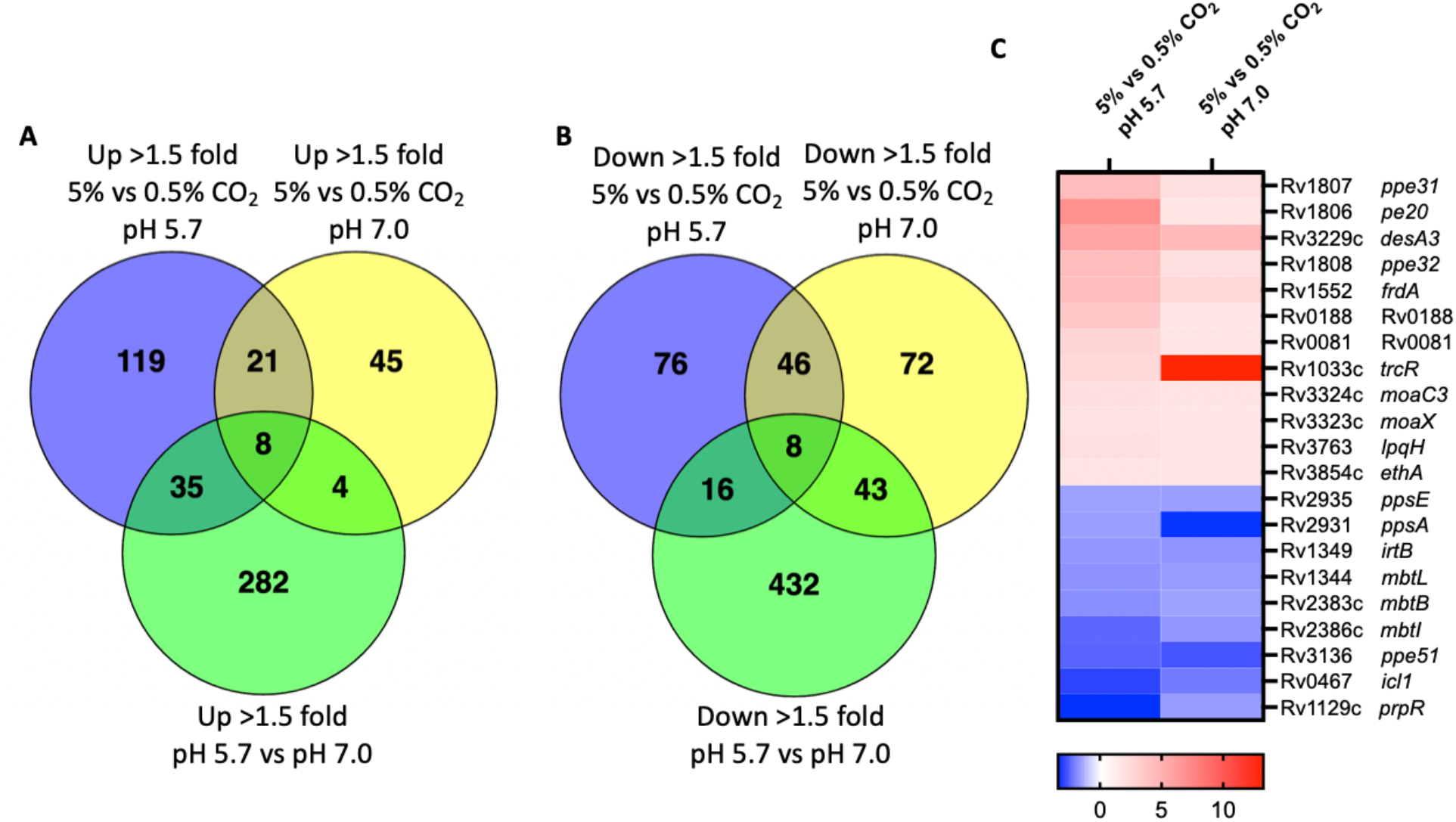
Significant overlap observed between expression profiles of increasing CO_2_ pressure at both pH 5.7 and pH 7.0. Venn diagrams comparing **A)** up-regulated or **B)** down-regulated genes (> 1.5 fold) modulated by 5% CO_2_ at pH 5.7 or 5% CO_2_ at pH 7.0 against the pH-induced or repressed regulon, respectively[11]. **C)** A heat map summarizing 21 of the 67 overlapping CO_2_-dependent, pH-independent regulated genes (>1.5-fold, q< 0.05). Induced genes (Up > 1.5-fold, 5% vs 0.5% CO_2_ at pH 5.7 and pH 7.0) are highlighted red and repressed genes (Down > 1.5-fold, 5% vs 0.5% CO_2_ at pH 5.7 and pH 7.0) are highlighted blue. Genes are annotated using the H37Rv genome.

For the CO_2_ regulated genes, independent of acidic pH, we observed 21 overlapping genes that were up-regulated by 5% CO_2_ at both pH 7.0 and 5.7 (Figure 5A and Table 2) and 46 overlapping genes that were down-regulated (Figure 5B and Table 3). Genes that responded to CO_2_, independent of pH, include intermediary metabolism genes (*icl1*,*prpR, frdA*), lipid metabolism genes (*desA3, ppsA, ppsE*), iron homeostasis genes (*irtB, mbtI, mbtL*), and hypoxia-induced genes (*Rv0081, Rv0188*) (Figure 5C, Table 2 and Table 3). Interestingly, we also see PE and PPE genes modulated that have been previously described and their functions resolved. These include PPE51, a nutrient transporter, and the PE20/PPE31 complex, which has been shown to mediate Mg^2+^ transport across the outer membrane[37]. PhoPR and acidic pH dependent adaptations have been linked to both PPE51 function [38, 39] and magnesium[6, 40], further linking CO_2_ to these pathogenesis associated pathways. Notably, we also observed the induction of *trcR*, the response regulator of the *trcRS* two-component regulatory system (TCS). Further analysis of all CO_2_ transcriptional profiles revealed a pattern of up-regulation or down-regulation of the *trcRS* TCS in response to changing CO_2_ levels (Table S2A-2E). The pattern of transcriptional changes in *trcRS* were independent of pH, however, *trcRS* is more highly induced overall by CO_2_ at pH 7.0 compared to pH 5.7. Interestingly, *trcR* is the most highly up-regulated gene with a fold induction of ~13-fold when comparing 5% CO_2_ to 0.5% CO_2_ at pH 7.0 (Table S2D), similarly *trcS* is the third most highly upregulated gene (4-fold induction). Interestingly, *canB*, showed an expression pattern similar to *trcRS*, with strong induction by CO_2_ at 5% vs. 0.5% CO_2_ at pH 7.0. These data demonstrate the existence of a CO_2_ regulon independent of pH and support the hypothesis that CO_2_ may be a putative signal for the *trcRS* TCS.

**Table 1.** Genes induced at 5% CO_2_ vs 0.5% CO_2_ (> 1.5 fold, q<0.05) at pH 5.7 and pH 7.0 as determined by Venn diagram overlap.

| **Rv Number** | **Gene Name** | **Fold Change (pH 5.7)** | **Fold Change (pH 7.0)** | **Description** |
| --- | --- | --- | --- | --- |
| MT3426 |  | 2.75 | 1.77 |  |
| MT3762 |  | 5.50 | 2.45 |  |
| Rv0077c | Rv0077c | 2.01 | 1.78 | Probable oxidoreductase |
| Rv0081 | Rv0081 | 2.75 | 1.71 | Predicted to response to early hypoxia responses, is a regulatory hub, transcriptional regulator (ArsR family) |
| Rv0188 | Rv0188 | 3.57 | 1.59 | Early hypoxia induced antigen, probable conserved transmembrane protein |
| Rv0458 | Rv0458 | 2.19 | 1.56 | Probable aldehyde dehydrogenase |
| Rv0459 | Rv0459 | 1.77 | 1.77 | Conserved hypothetical protein |
| Rv1033c | *trcR* | 2.46 | 13.24 | Two-component response regulator |
| Rv1552 | *frdA* | 4.15 | 2.57 | Fumarate reductase flavoprotein subunit |
| Rv1806 | *pe20* | 6.95 | 1.72 | PE-family protein |
| Rv1807 | *ppe31* | 7.77 | 3.17 | PPE-family protein |
| Rv1808 | *ppe32* | 4.30 | 2.13 | PPE-family protein |
| Rv1926c | *mpt63* | 3.14 | 1.59 | Immunogenic protein |
| Rv2557 | Rv2557 | 9.85 | 1.77 | Conserved hypothetical protein |
| Rv2558 | Rv2558 | 5.11 | 1.50 | Conserved hypothetical protein |
| Rv3196A | Rv3196A | 1.51 | 1.57 | Hypothetical protein |
| Rv3229c | *desA3* | 5.64 | 4.42 | Possible linoleoyl-CoA desaturase |
| Rv3230c | Rv3230c | 1.82 | 2.44 | Hypothetical oxidoreductase |
| Rv3323c | *moaX* | 1.90 | 1.59 | Probable MoaD-MoaE fusion protein MoaX |
| Rv3324c | *moaC3* | 2.17 | 1.73 | Molybdenum cofactor biosynthesis, protein C |
| Rv3854c | *ethA* | 1.77 | 1.71 | Monooxygenase/ activates prodrug ethionamide |

**Table 2.**
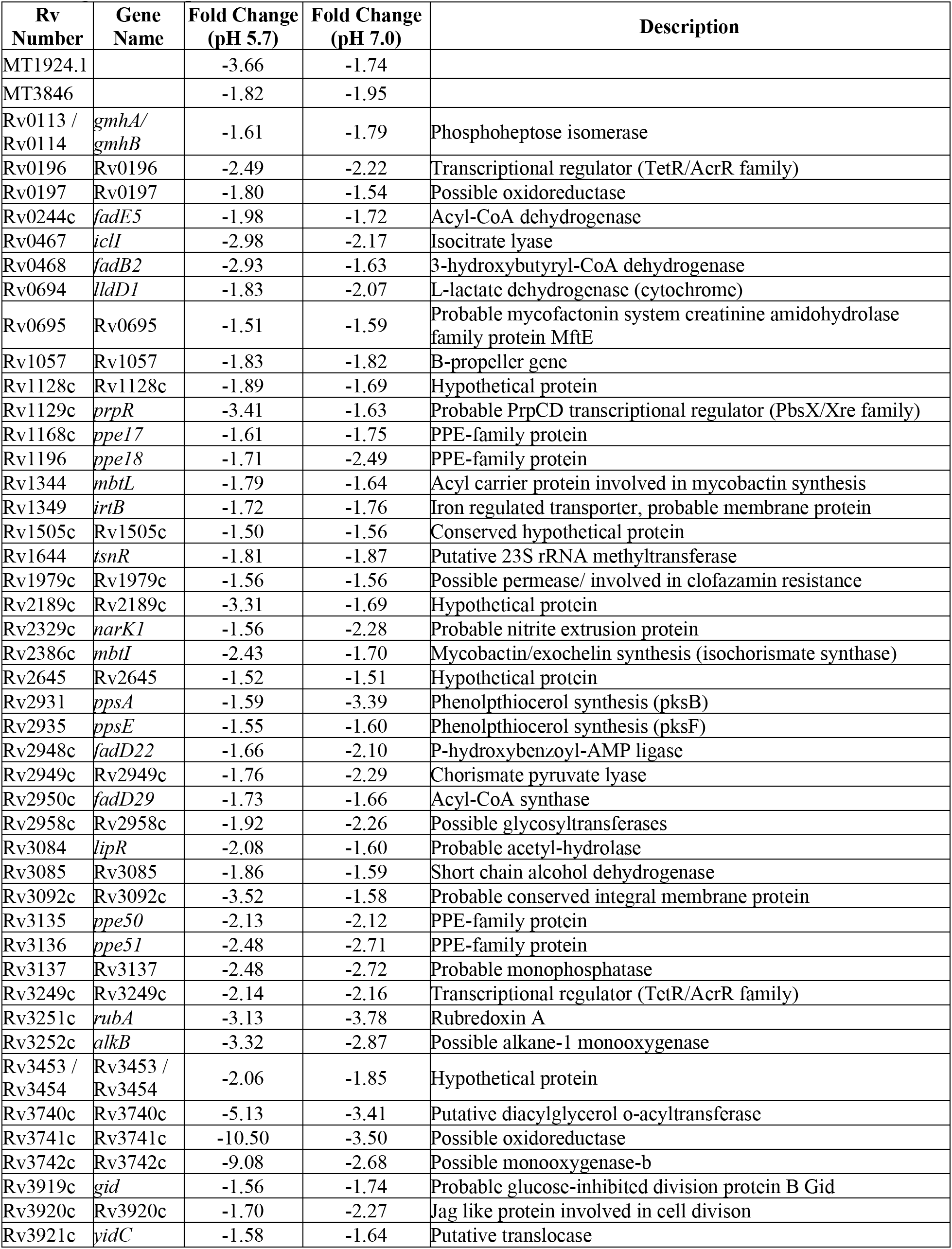
Genes repressed at 5% CO_2_ vs 0.5% CO_2_ (> 1.5 fold, q<0.05) at pH 5.7 and pH 7.0 as determined by Venn diagram overlap.

**Table 3.**
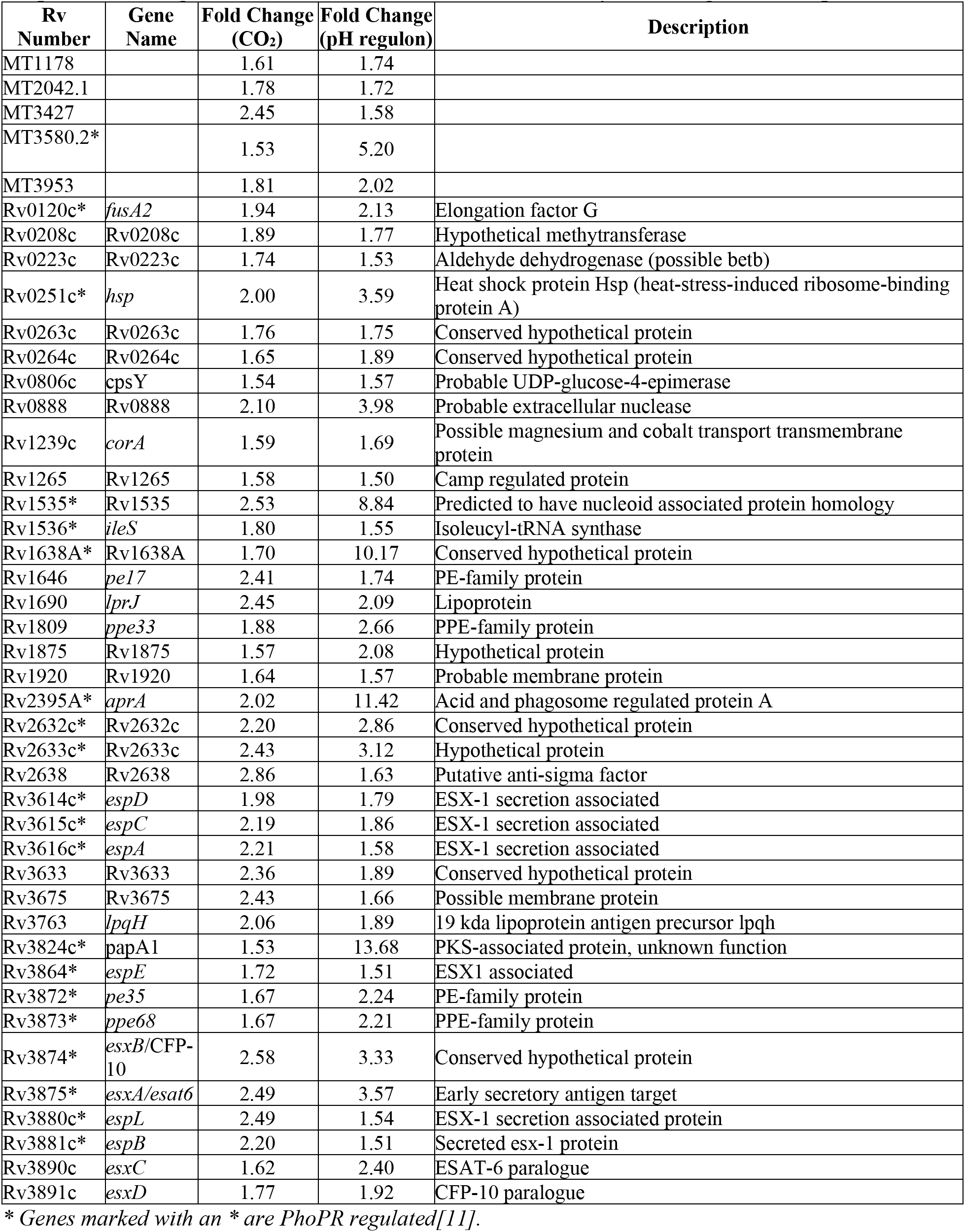
Genes induced at 5% CO_2_ vs 0.5% CO_2_ at pH 5.7 (> 1.5 fold, q<0.05) compared to genes in the pH-induced regulon (> 1.5 fold, q<0.05)[11] as determined by Venn diagram overlap.

## Discussion

Here, we have demonstrated that increasing CO_2_ concentration induces PhoPR signaling, and its induction is independent of medium pH (Figure 1A and B, Table S2A-E). These finding are consistent with PhoPR acting as a CO_2_ sensor. During infection, Mtb can encounter a variety of changing CO_2_ conditions that might be important for PhoPR-dependent virulence. For example, as Mtb infects humans from respiratory droplets or aerosols in the environment (low environmental CO_2_ concentrations) into the host lung environment (moderate, physiological CO_2_ concentrations), CO_2_ could provide a key cue that the bacterium has entered the host. PhoPR and high CO_2_ is required for key steps of initial macrophage infection (*e.g*. ESX-1 secretion), therefore, inducing these pathways at the onset of infection, prior to macrophage infection, could enhance virulence. PhoPR also induces the synthesis of sulfolipid, a cell envelope lipid that causes animals to cough [9]. As TB infection progresses lung damage can obstruct airflow and cause hypercapnic environments[41, 42]. These high levels of CO_2_ could trigger Mtb to generate more sulfolipid and drive the cough response and transmission. As such, CO_2_ may play a critical role in both inducing signaling cascades for survival during macrophage infection and transmission to new hosts. Thus, studying CO_2_ as an environmental cue may provide important new insights into Mtb pathogenesis.

PhoPR is strongly induced by acidification of the host macrophage early in infection[1]. We hypothesized that the proton produced during the catalysis of CO_2_ hydration by CA is a possible mechanism linking CO_2_ and PhoPR[15]. This model is further supported by the carbonic anhydrase inhibitor, ETZ, modulating the PhoPR regulon. Using knockdowns or knockouts of the three CA, we found that only *canB* was required for virulence. We reasoned that since both *canB* and *phoPR* are required for virulence in macrophages, that the virulence phenotype of *canB* knockdown may be related to loss of PhoPR signaling and that *canB* is required for PhoPR induction. However, under conditions that stimulate PhoPR signaling (5% CO_2_ and pH 5.7), we observed no impact on *aprA* regulation (a PhoPR signaling biomarker) in the *canB* knockdown. Thus, the link between CA, CO_2_ and PhoPR signaling remains unresolved. Additionally, the mechanism by which *canB* has reduced virulence in macrophages is similarly unresolved. It is possible that residual CA activity present in the CRISPRi knockdowns is sufficient to promote PhoPR signaling at acidic pH. Indeed, our prior study of ETZ in whole cell Mtb, shows ETZ completely inhibits CA activity at the concentrations tested[11]. CAs are efficient enzymes, so residual activity is a plausible explanation, even with a 100-fold reduction of CA expression in the knockdowns. Knockout strains in *canA, canB* or in combination would be needed to test this hypothesis. However, efforts to generate a *canB* knockout have been unsuccessful, despite evidence it is non-essential[43]. Therefore, further experiments are needed to definitively refute the hypothesis that CO_2_ modulates PhoPR signaling through a CA-dependent mechanism, although the data presented here

This work has provided a foundation to begin dissecting the link between PhoPR signaling, CA activity, and sensing changes in environmental CO_2_. We propose a hypothetical model where changes of CO_2_ do not change the pH of the extracellular medium because of the high levels of buffers in the medium. However, the buffer may not permeate the mycomembrane, resulting in localized changes in pH in the pseudoperiplasm that differ from the extracellular medium. In this model, high CO_2_ permeates through the mycomembrane and results in enhanced accumulation of protons in the pseudoperiplasm due to the CA-dependent or -independent conversion of CO_2_ to bicarbonate and a proton. Alternatively, it is possible that PhoPR is responding to shared changes in metabolism or cell envelope driven by CO_2_ and acidic pH, such as changes to the methylcitrate cycle or PDIM synthesis genes, and that PhoPR is responding indirectly to the environmental signals. Identifying the biochemical mechanism of PhoPR activation is required to resolve models that support direct sensing of environmental cues as compared to indirectly sensing adaptations driven by environmental cues. Given the importance of PhoPR in Mtb pathogenesis, it is possible that PhoPR can respond to multiple environmental cues.

The specific induction of the TrcRS TCS in response to changing CO_2_ levels is interesting because it may indicate a previously unknown CO_2_-dependent signaling pathway. *trcR* encodes for the response regulator which is located directly upstream of the sensor kinase, *trcS*[44] [45]. While little is known about the conditions under which *trcRS* may be expressed, it is induced during early to mid-logarithmic growth phase under aerobic conditions *in vitro* and following initial macrophage infection at 18 hours but not after 48 hours[44]. Additionally, only one member of the TrcRS regulon has been defined, Rv1057. Rv1057 is a β propeller protein of unknown function, and its expression is repressed by TrcR[46, 47]. Interestingly, TrcRS is not the only regulator of Rv1057, which is also directly regulated by MprAB, another TCS that is associated with cell envelope stress[45, 46]. Similarly, loss of *mtrB* in *Mycobacterium smegmatis* leads to defects in cell morphology and cell division, which can be reversed by *trcS* overexpression. Based on these results and ours, it is possible that TrcRS is responding to changes in CO_2_ to modulate Mtb metabolism or the cell envelope.

When defining genes regulated by CO_2_ independent of pH, we found widespread differential expression of genes involved in intermediary metabolism and respiration or lipid metabolism. For example, genes induced by CO_2_ in a pH-independent manner include *ethA, moaX, moaC3, frdA, Rv3230c*, and *desA3* (Table 2). *moaX* and *moaC3* are involved in molybdenum cofactor biosynthesis which is required for oxidoreductase and nitrate reductase function[48]. *desA3* and the oxidoreductase, *Rv3230c*, interact to produce oleic acid which is essential for mycobacterial membrane phospholipids and triglycerides[49]. *desA3* is also essential for Mtb survival during infection[30]. We also see that half of CO_2_-repressed genes are involved in metabolic processes (Table 3) Notably, these include methylcitrate genes (*icl1, prpR*), phthiocerol dimycocerosate (PDIM) biosynthesis (*ppsAE, fadD22*, and *fadD29*), and hydrocarbon degradation (*Rv3249c, rubA, alkB*). Together, these expression changes indicate that CO_2_ induces metabolic shifts that could be required for its survival in the host. Indeed, Beste *et al*., have shown that Mtb fixes CO_2_ and does so using *icl1-*associated pathways[50]. One interesting observation is the induction of the *pe20-ppe33* locus by CO_2_ (Figure 5A and C, Tables 2 and 4). We see *pe20, ppe31*, and *ppe32* induced in a CO_2_-dependent, pH-independent manner while *ppe33* is induced both by CO_2_ and pH. This locus has been shown to be upregulated during Mg^2+^ starvation, clusters with the Mg^2+^ transporter *mgtC*, and possibly play a role in magnesium homeostasis[6, 51, 52]. Wang and colleagues recently showed that knockout mutants of this locus exhibit a growth defect in Mg^2+^-limiting media, especially at mildly acidic pH[37]. Likewise, Piddington *et. al*. demonstrated that Mtb requires higher levels of Mg^2+^ for growth at acidic pH[40]. The phagosome is thought to be a Mg^2+^-limiting environment[53]. Induction of this locus, specifically at higher CO_2_ levels, supports Mtb may be sensing the higher CO_2_ in the lungs and adapting its physiology accordingly for the nutrient-limiting environment of the alveolar macrophages, possibly priming itself for survival during infection. In support of this, we see overlap with published TrcR ChIP-Seq data [54] and CO_2_ RNA-Seq, notably strong co-regulation of *trcRS* and its binding targets, *desA3, ppe32*, Rv0458, and Rv3230c. These data support a model where CO_2_, via TrcR, induces metabolic changes in Mtb that prime Mtb for the nutrient-restricted environment of the host phagosome.

**Table 4.** Genes repressed at 5% CO_2_ vs 0.5% CO_2_ at pH 5.7 (> 1.5 fold, q<0.05) compared to genes in the pH-repressed regulon (> 1.5 fold, p<0.05) [11] as determined by Venn diagram overlap.

| Rv Number | Gene Name | Fold Change (CO<sub>2</sub>) | Fold Change (pH regulon) | Description |
| --- | --- | --- | --- | --- |
| MT0600 |  | -1.78 | -2.58 |  |
| MT1775 |  | -1.54 | -2.26 |  |
| MT2617 |  | -1.75 | -1.66 |  |
| Rv0972c | *fadE12* | -1.57 | -3.25 | Acyl-CoA dehydrogenase |
| Rv1169c | *lipX* | -1.76 | -1.73 | Possible lipase/ PE-family protein |
| Rv1195 | *PE13* | -2.79 | -2.30 | PE-family protein |
| Rv1297 | *rho* | -1.78 | -1.73 | Transcription termination factor rho |
| Rv1733c | Rv1733c | -2.05 | -2.10 | Probable conserved transmembrane protein |
| Rv1737c | *narK2* | -1.77 | -3.38 | Nitrite extrusion protein |
| Rv1738 | Rv1738 | -1.56 | -3.90 | Possibly interact with ribosome structural prection |
| Rv2007c | *fdxA* | -2.08 | -2.49 | Ferredoxin |
| Rv2028c | Rv2028c | -2.54 | -1.86 | Universal stress protein family protein |
| Rv2029c | *pfkB* | -1.90 | -3.06 | Phosphofructokinase |
| Rv2030c | Rv2030c | -1.98 | -2.47 | Conserved protein |
| Rv2031c | *hspX* | -2.20 | -1.93 | Heat shock protein |
| Rv2032 | *acg* | -1.54 | -2.31 | Conserved protein |
| Rv2379c | *mbtF* | -1.54 | -1.56 | Mycobactin/exochelin synthesis (lysine ligation) |
| Rv2382c | *mbtC* | -1.79 | -1.51 | Mycobactin/exochelin synthesis |
| Rv2383c | *mbtB* | -1.83 | -1.50 | Mycobactin/exochelin synthesis (serine/threonine |
| Rv2450c | *rpfE* | -2.61 | -1.84 | Probable resuscitation promoting factor |
| Rv2987c | *leuD* | -2.03 | -2.56 | 3-isopropylmalate dehydratase small subunit |
| Rv2988c | *leuC* | -2.92 | -1.96 | 3-isopropylmalate dehydratase large subunit |
| Rv2989 | Rv2989 | -2.20 | -1.70 | Transcriptional regulator (iclR family) |
| Rv3402c | Rv3402c | -2.11 | -1.67 | Conserved protein |

In conclusion, we report here experimental evidence that defines a link between CO_2_ levels and PhoPR signaling. The impetus of this study was to define the mechanism by which the CA inhibitor ETZ inhibits PhoPR signaling. However, the overarching hypothesis driving this study, that CO_2_ regulates PhoPR in an CA dependent manner, remains unresolved. It is possible that the CRISPRi knockdown of *canAB* is not sufficient to elicit a change in PhoPR signaling and that CA knockouts are required to replicate the ETZ-dependent inhibition of PhoPR signaling. There may also be additional CA not annotated in the Mtb genome that ETZ could be inhibiting. Or, potentially, ETZ could be targeting something else altogether, even directly inhibiting PhoPR. Further studies will be required to resolve these questions. Nevertheless, important new discoveries have resulted from these studies. Our findings show that PhoPR functions as a CO_2_ sensor, including regulation of the central regulator of Mtb virulence, the ESX-1 system, by CO_2_. We also found that increasing CO_2_ concentrations elicit a core CO_2_-expression profile and that includes changes in metabolism and regulation of the TrcRS regulon. In addition, *canB* is shown to be required for virulence in macrophages, while *canA* and *canC* are dispensable. Thus, we have defined a complex interplay between CO_2_, acidic pH and PhoPR in regulating Mtb gene expression and virulence that supports further investigation of the mechanisms linking these physiologies and defining their role in pathogenesis.

## Acknowledgements

We thank Sarah Fortune and Jeremy Rock for sharing the CRISPRi plasmids and Christopher Sassetti for sharing the ORBIT plasmids used in this study. We also thank the MSU RTSF for technical support for the RNA-seq library preparation and sequencing and the MSU Flow Cytometry Core for assistance with the flow cytometry analysis and sorting. This work was supported by a grant from the NIH NIAID (R01AI116605) and AgBioResearch.

## Author Contributions

S.J.D., B.K.J., R.G., and R.B.A. conceived the project. R.G. and B.K.J. performed the fluorescence reporter experiments. S.J.D. carried out the RNA-seq analysis, qRT-PCR studies, and macrophage infections. R.G. and S.J.D. constructed and confirmed the CRISPRi constructs and ORBIT knockouts. R.G. conducted the RNAseq experiments. S.J.D. and R.B.A. wrote the manuscript.

---

**Supplemental Figure 1.**
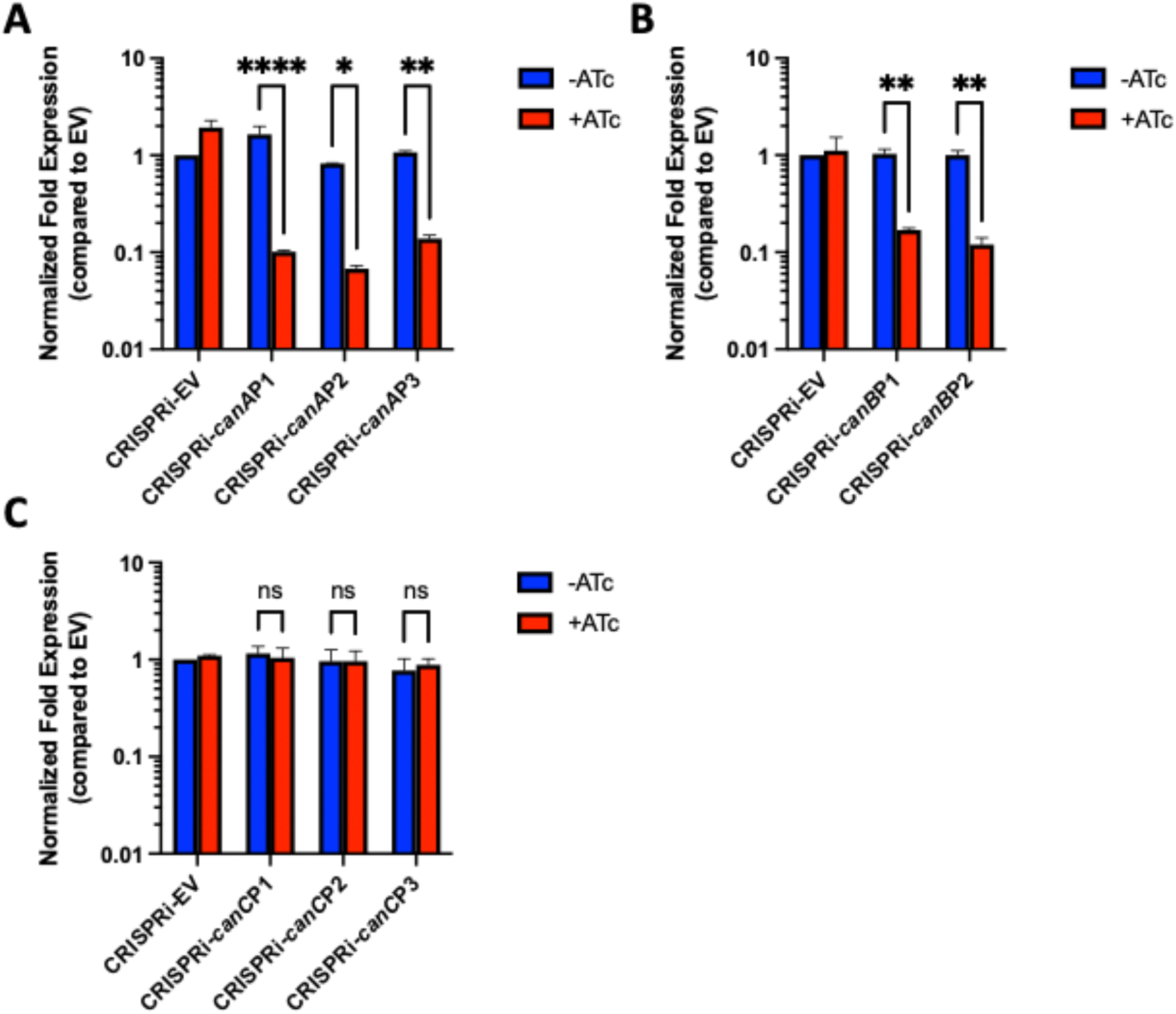
qRT-PCR confirmation of *canA* and *canB* CRISPRi in WT CDC1551. **A)** dCas9_Sth1_ knockdown of *canA* target in Mtb. Three sgRNAs targeting *canA* were co-expressed with dCas9_Sth1_ (+ATc). After 6 days of incubation in 7H9 media, total RNA was extracted, and *canA* knockdown was quantified by qRT-PCR. **B)** Two sgRNAs targeting *canB* were co-expressed with dCas9_Sth1_ (+ATc) and *canB* knockdown was quantified by qRT-PCR. **C)** Three sgRNAs targeting *canC* were co-expressed with dCas9_Sth1_ (+ATc). *canC* knockdown was quantified by qRT-PCR but lacked knockdown efficiency. Error bars in all three figures represent the standard deviation of three technical replicates. Significance was determined by two-way ANOVA (Šídák’s multiple comparisons test; ****P < 0.0001). Mean ± SD are shown in the bar graph.

**Supplemental Figure 2.**
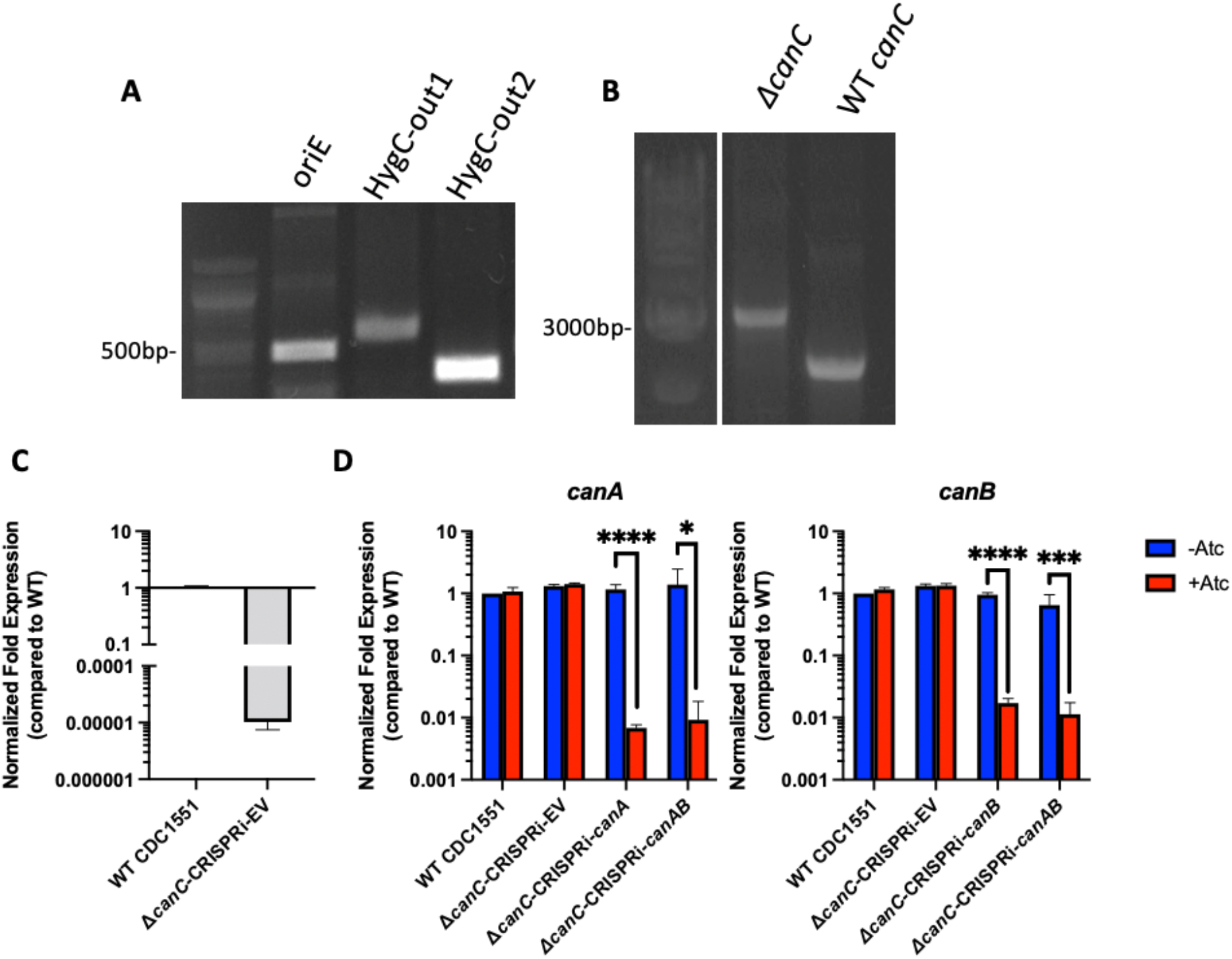
PCR and qRT-PCR confirmation of *canC* ORBIT knockout and CRISPRi. **A)** PCR amplification of the 5’(oriE) and 3’ (HygC-out1/2) junctions of CDC1551 Δ*canC*. **B)** PCR analysis of the integration site of the payload plasmid (pKM464). pKM464 is 3082 bp which is consistent with the size of the bands observed in the *canC* deletion mutants compared to WT *canC* which is 2295 bp. **C)** qRT-PCR analysis confirming the *canC* knockout. Total RNA was collected after samples were grown for six days in 7H9 media buffered to pH 7.0. Fold expression was normalized to WT. Error bars represent the standard deviation of three technical replicates. Deletion mutant of *canC* typically exhibited a non-specific primed Ct ~35 cycles compared to WT which had a Ct of ~16 cycles. **D)** qRT-PCR confirmed gene knockdown of *canA, canB,and canAB* in the *canC* deletion mutation background. Fold expression was normalized to WT Mtb. Error bars represent the standard deviation of three technical replicates. Significance was determined by student’s t-test. (*P < 0.05, ***P < 0.001, ****P < 0.0001).

**Supplemental Figure 3.**
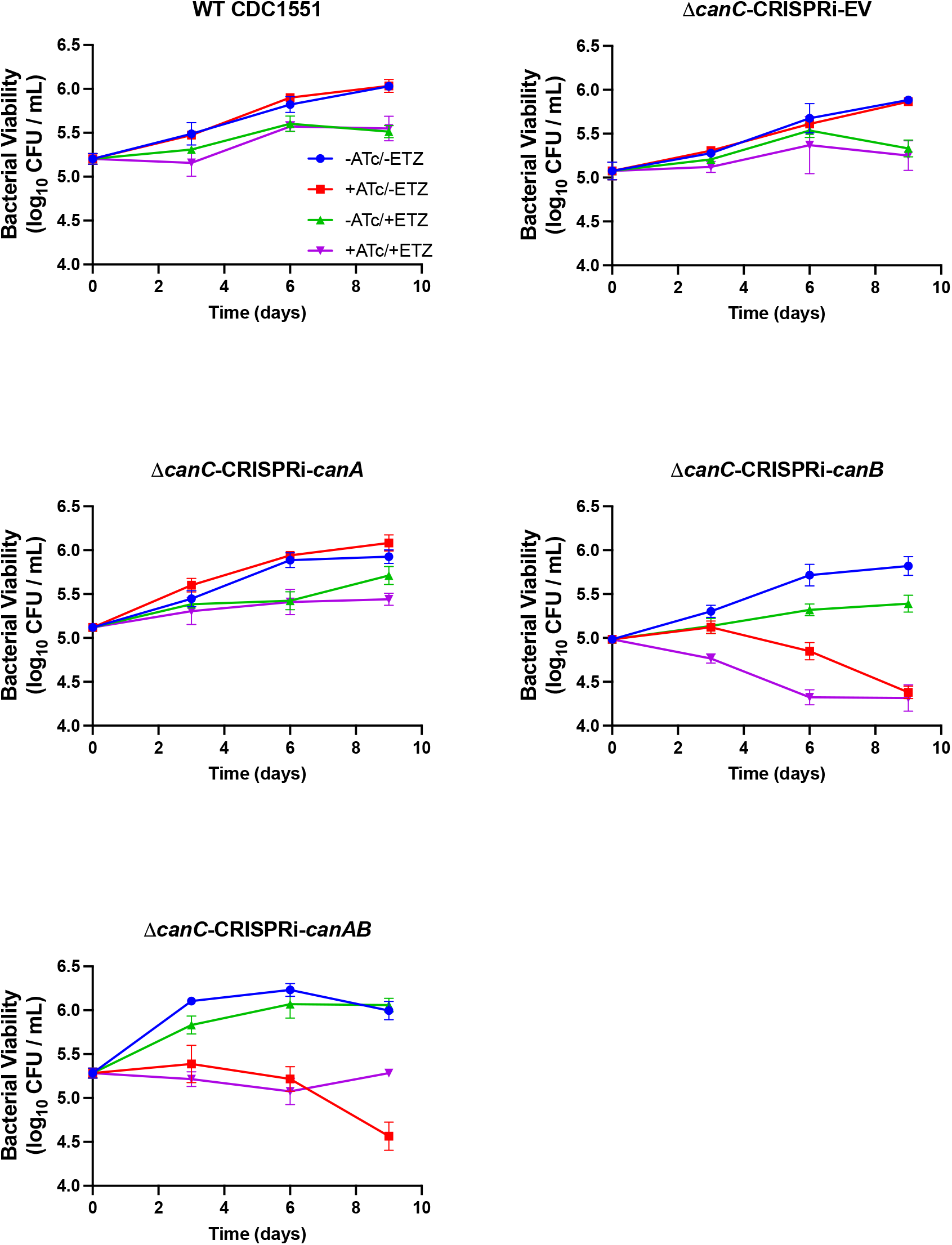
Nine day bacterial viability CFUs that correspond to the endpoint data summarized in Figure 2E. BMDMs infected with the CA CRISPRi strains in the CDC1551 Δ*canC* background. All strain treatments were performed in triplicate. Error bars indicate standard deviation.

**Supplemental Figure 4.**
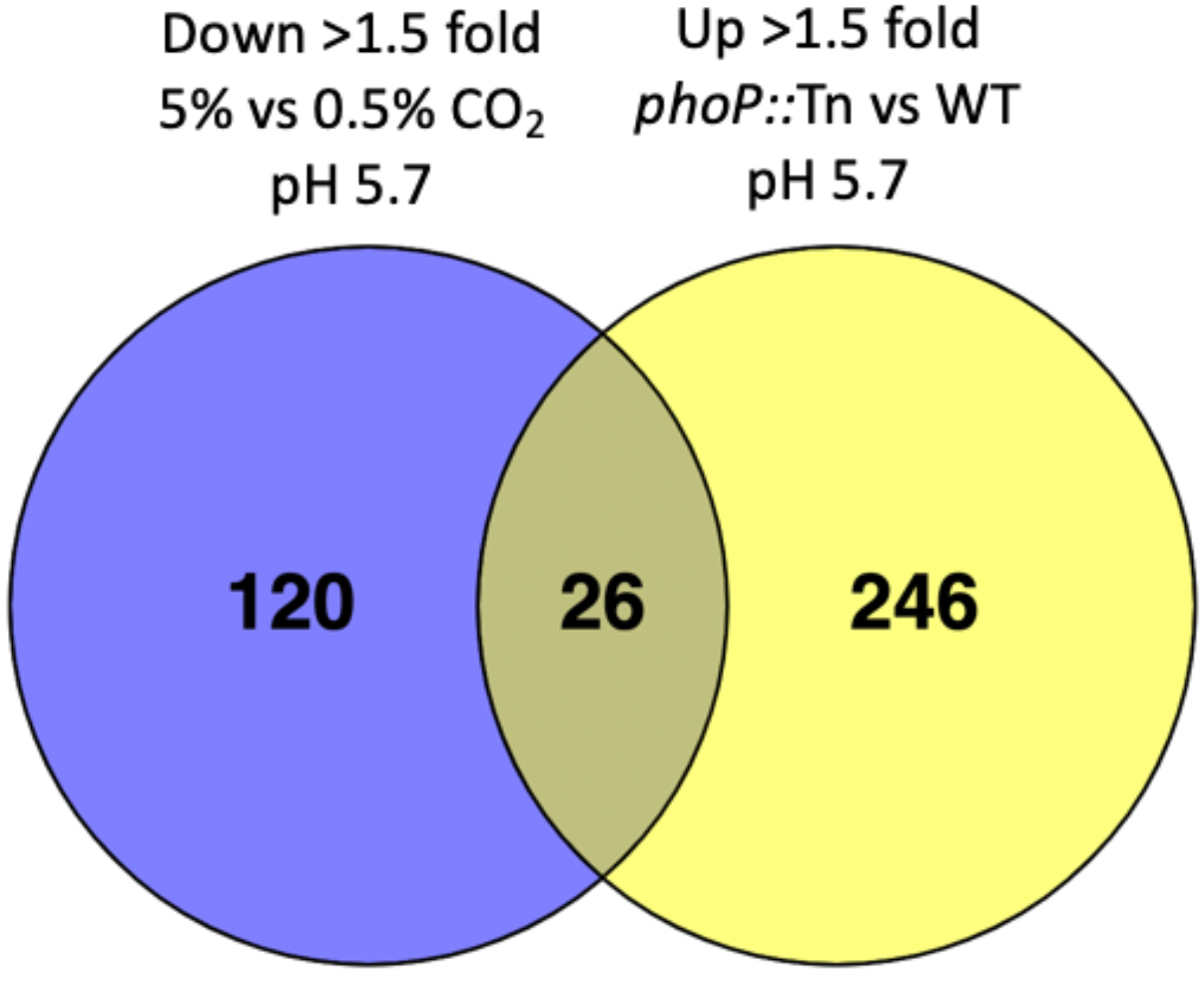
Venn diagram of down-regulated genes 5% CO_2_ vs 0.5% CO_2_, pH 5.7 compared to up-regulated *phoP::Tn* profile. Significant gene overlap observed between genes downregulated (Down) (>1.5-fold, q < 0.05) by 5% CO_2_ treatment at pH 5.7 and Upregulated (Up) (>1.5-fold, q < 0.05) in the *phoP::Tn* mutant strain at pH 5.7[11]

**Table S1.**
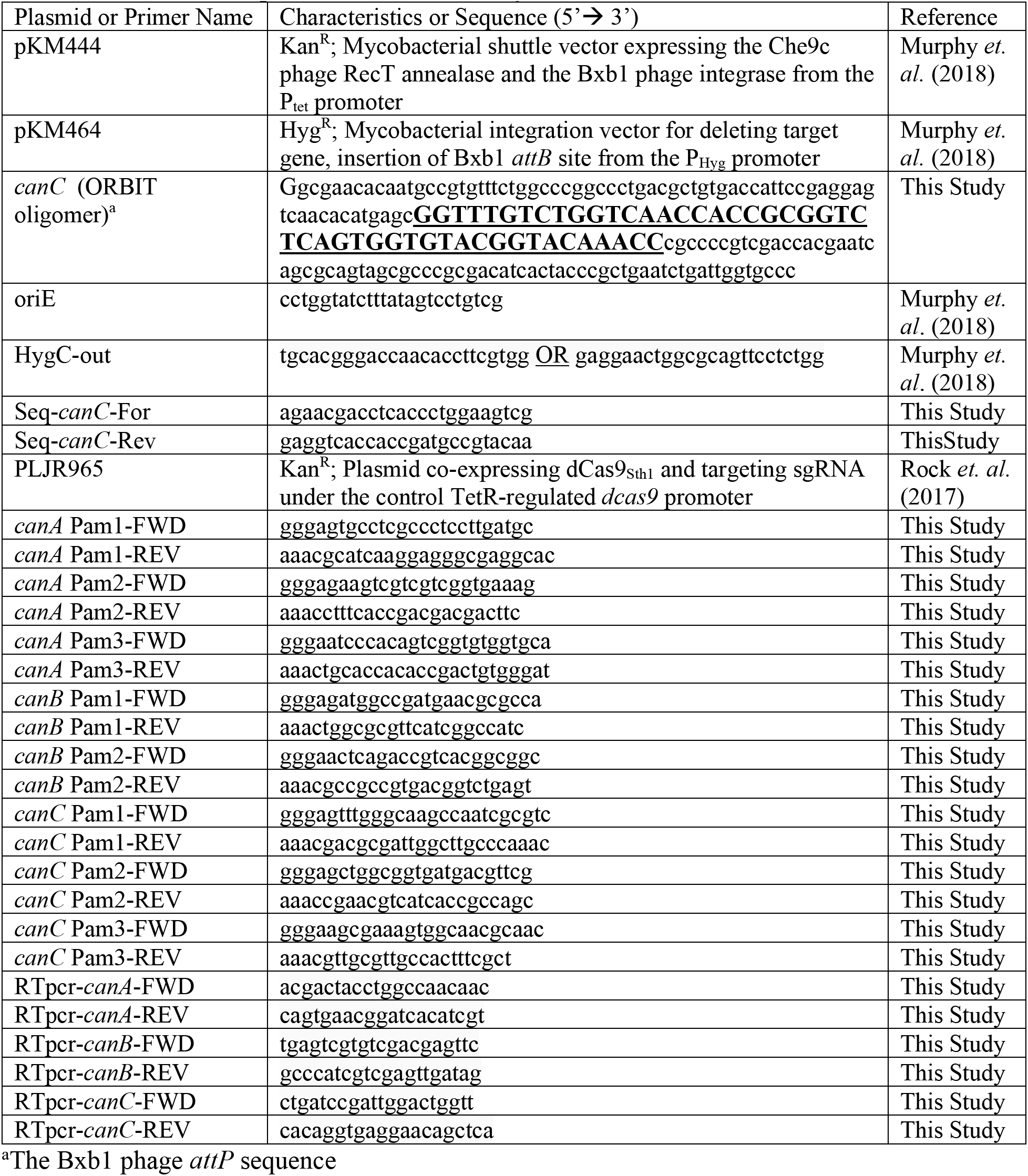
Plasmids and primers used in this study.

